# Estimation of in-scanner head pose changes during structural MRI using a convolutional neural network trained on eye tracker video

**DOI:** 10.1101/2021.03.04.433975

**Authors:** Heath R Pardoe, Samantha P Martin, Yijun Zhao, Allan George, Hui Yuan, Jingjie Zhou, Wei Liu, Orrin Devinsky

**Author notes:** Corresponding author: Heath Pardoe, PhD, Department of Neurology, NYU Grossman School of Medicine 145 East 32^nd^ St, 5^th^ Floor, room 507, New York City, NY 10016, USA.

## Abstract

**Introduction:** In-scanner head motion is a common cause of reduced image quality in neuroimaging, and causes systematic brain-wide changes in cortical thickness and volumetric estimates derived from structural MRI scans. There are few widely available methods for measuring head motion during structural MRI. Here, we train a deep learning predictive model to estimate changes in head pose using video obtained from an in-scanner eye tracker during an EPI-BOLD acquisition with participants undertaking deliberate in-scanner head movements. The predictive model was used to estimate head pose changes during structural MRI scans, and correlated with cortical thickness and subcortical volume estimates.

**Methods:** 21 healthy controls (age 32 ± 13 years, 11 female) were studied. Participants carried out a series of stereotyped prompted in-scanner head motions during acquisition of an EPI-BOLD sequence with simultaneous recording of eye tracker video. Motion-affected and motion-free whole brain T1-weighted MRI were also obtained. Image coregistration was used to estimate changes in head pose over the duration of the EPI-BOLD scan, and used to train a predictive model to estimate head pose changes from the video data. Model performance was quantified by assessing the coefficient of determination (R^2^). We evaluated the utility of our technique by assessing the relationship between video-based head pose changes during structural MRI and (i) vertex-wise cortical thickness and (ii) subcortical volume estimates.

**Results:** Video-based head pose estimates were significantly correlated with ground truth head pose changes estimated from EPI-BOLD imaging in a hold-out dataset. We observed a general brain-wide overall reduction in cortical thickness with increased head motion, with some isolated regions showing increased cortical thickness estimates with increased motion. Subcortical volumes were generally reduced in motion affected scans.

**Conclusions:** We trained a predictive model to estimate changes in head pose during structural MRI scans using in-scanner eye tracker video. The method is independent of individual image acquisition parameters and does not require markers to be to be fixed to the patient, suggesting it may be well suited to clinical imaging and research environments. Head pose changes estimated using our approach can be used as covariates for morphometric image analyses to improve the neurobiological validity of structural imaging studies of brain development and disease.

## 1. Introduction

Head motion is a fundamental source of error in MRI. Recent studies have demonstrated that subtle in-scanner head motion is associated with systematic brain-wide underestimation of quantitative cortical thickness and regional volume estimates derived from structural MRI [1–4]. Because in-scanner head motion is increased in children, the elderly, and in clinical populations relative to healthy controls [2, 4], these systematic effects may cause false positive findings in studies that use MRI-based morphometric estimates to characterize differences between study populations. In-scanner motion also contributes to within group variability in morphometric estimates [5], which reduces statistical power and increases the likelihood of false negative findings.

Here, we trained a convolutional neural network (CNN) to estimate changes in head pose over the duration of an image acquisition using in-scanner video obtained from an MRI compatible eye tracker. These head pose estimates were then used as predictor variables in statistical analyses of cortical thickness and volumes of subcortical structures derived from structural MRI. Ground truth head pose changes were obtained by acquiring video data during an echo-planar imaging (EPI) acquisition typically used for functional MRI (fMRI) experiments, with study participants undertaking prompted, deliberate head motions during the scan. Standard coregistration methods typically used for motion correction in fMRI studies were used to estimate changes in head pose from the EPI acquisition. The model was then applied to video obtained during structural image acquisitions. To test the utility of our approach, we applied the predictive model to video obtained during whole brain T1-weighted MRI scans obtained with subjects (i) holding their head still, as per a standard image acquisition, and (ii) deliberately moving their head using prompted stereotypical movements during the acquisition in order to obtain motion-affected MRI scans. We used these structural MRI scans to derive whole brain cortical thickness maps and subcortical volume estimates, and carried out statistical analyses to determine if measured cortical thickness and subcortical volumes were significantly correlated with video-based head pose estimates. Prior studies that inferred in-scanner motion using proxy methods [2–4] suggested a brain-wide reduction in cortical thickness and volume estimates with increased head motion. Our proposed technique allows for non-invasive, acquisition agnostic assessment of head pose changes at a high temporal frequency (30 Hz).

The problem of head pose estimation from video is well established in the computer vision literature [6]. Our specific use case is challenging because the eye tracker video field of view is limited to the upper facial region encompassing a single eye (Figure 1). Parts of the face are also occluded by the MRI coil hardware. Nevertheless convolutional neural networks have performed well in a number of head pose estimation problems, motivating their use in our study [7].

**Figure 1.**
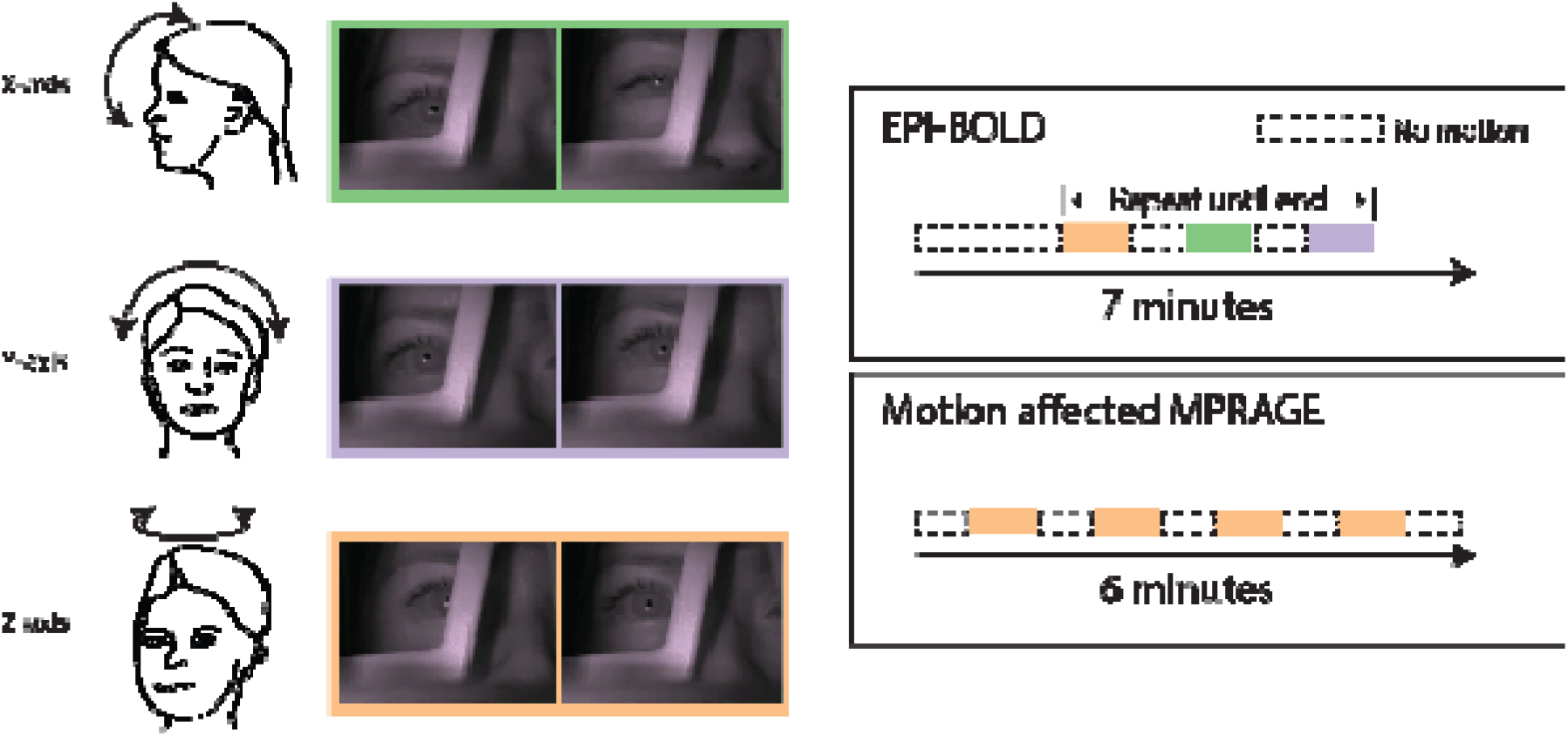
Overview of deliberate in-scanner head motions used in our study. For the EPI-BOLD acquisition, the head was held still for the first 2 minutes of the scan then repeating head motions were carried out interleaved with 30 second no-motion epochs until the end of the scan. During the motion affected image acquisition the head was rotated around the z-axis (“no-no” motion) interleaved with 30 second no-motion epochs.

Various methods have been proposed to detect and estimate motion in MRI, primarily with the goal of obtaining images free from motion-related artifacts. These methods include prospective techniques that modify image acquisition parameters during scanning to reduce motion artifacts, and retrospective techniques that assess motion and correct for motion-related effects after imaging data was obtained [5, 8–10]. Recent studies have utilized external optical signals, electromagnetic motion trackers or short volumetric navigators interleaved with the neuroanatomical image acquisition sequence to estimate and correct for changes in head pose during imaging [5, 10–18]. To the best of our knowledge these techniques are not currently widely used in a clinical imaging setting, however recent large-scale multi-center imaging studies such as the Adolescent Brain Cognitive Development (ABCD) study and the Human Connectome Project (HCP) Lifespan Development and Aging projects utilize volumetric navigators in their neuroanatomical imaging sequences to correct for in-scanner head motion [10, 17, 19, 20]. For the HCP Lifespan Development and Aging studies less than 5% of neuroanatomical scans have been rated as poor quality; the investigators have attributed this low number of poor quality scans to the use of these motion-robust sequences [19]. A modest number of prior studies have directly estimated in-scanner head motion during scanning and compared quantitative estimates of neuroanatomical properties such as cortical thickness or volumes of brain structures [1, 5, 18]. Our proposed method may be a practical approach as a quality assurance tool for estimating head motion during MRI acquisition because eye trackers are widely available in clinical research centers, do not require placing markers on the study participant, and do not require modification of imaging sequences.

Specific hypotheses investigated in this study were:

a. Head pose estimates derived from our CNN-based head pose prediction model would be correlated with fMRI-based head pose estimates in an independent hold-out dataset of subjects not used to train the model.
b. Cortical thickness and subcortical volume estimates derived from motion-affected and motion-free neuroanatomical image acquisitions would be correlated with head pose changes estimated using our CNN-based head pose prediction model.

## 2. Methods

To facilitate further methods development we have provided links to imaging data, video files, and code associated with the study at the following site: https://sites.google.com/site/hpardoe/headmotion

### 2.1 Participants

Twenty one healthy adult participants were recruited from the community for this study (age 32 ± 13 years, 11 female). Approval for the study was obtained from the NYU Langone Health Institutional Review Board, and written informed consent to participate in the study was obtained from all study participants.

### 2.2 Imaging

Imaging was obtained on a 3T Siemens Prisma magnet equipped with an Eyelink 1000 Plus in-scanner eye tracker. Video was recorded using a USB video capture card connected to the eye tracker monitor. The imaging protocol consisted of four whole brain T1-weighted MPRAGE acquisitions: (i) two motion-affected MPRAGE scans and (ii) two motion-free MPRAGE scans, and a seven minute EPI-BOLD acquisition. The image acquisition order was: (1) Motion-free MPRAGE, (2) Motion-affected MPRAGE, (3) EPI-BOLD acquisition, (4) Motion-affected MPRAGE, (5) Motion-free MPRAGE. Image acquisition parameters were:

Whole brain T1-weighted MPRAGE: 1mm isotropic voxel size, echo time (TE) = 3 ms, flip angle = 8 degrees, inversion time (TI) = 0.9 s, repetition time = 2.5 s.

EPI-BOLD: 2 mm isotropic voxel size, 324 volumes acquired, repetition time = 1.3 s, echo time (TE) = 35 ms, multiband acceleration factor = 4, total acquisition time = 7 minutes.

In-scanner head motions were carried out in a reproducible manner by presenting a video demonstrating the required head movements that was viewed by the participant during the scan. The video included timing information to indicate when the next head motion was due to occur. Figure 1 provides a schematic overview of the study experimental paradigm. The EPI-BOLD head motion paradigm consisted of (i) A head nodding motion similar to the head motion indicating “yes”, primarily consisting of rotation about the x-axis in the participant’s plane of reference, (ii) A head turning motion similar to the head motion indicating “no” (rotation around the z-axis), and (iii) A side-to-side head motion with ears moving from shoulder to shoulder (rotation around the y-axis). Each of these deliberate motions took approximately 20 seconds with a 30 second break between motions. For each type of movement, the head was held at the maximal values for ~5-6 seconds to ensure the EPI-BOLD acquisition captured image volumes at the extremes of each type of deliberate head motion.

For the motion-affected structural MRI scans, we instructed participants to only undertake the “no” head motion as described in (ii) above. Each “no” motion was interspersed with a 30 second break. The “no” motion is effectively a rotation around the long axis of the MRI scanner which corresponds to the z-axis for head pose analyses.

Head pose changes over the duration of the EPI-BOLD acquisition were estimated using the program “MCFLIRT” (Motion Correction FMRIB Linear Image Registration Tool), provided as part of the FSL software (https://fsl.fmrib.ox.ac.uk/fsl/fslwiki/MCFLIRT, [21]). In brief, we used MCFLIRT to carry out sequential rigid-body coregistration of volumes in the EPI-acquisition to the first volume acquired in the acquisition. The six parameters describing the coregistration (3 rotation and 3 translation parameters) were used as target predictor variables to train the video-based predictive model.

Head pose changes during EPI-BOLD imaging in our participants were compared with head pose changes estimated from resting state fMRI acquisitions in the ABIDE cohort [22]. This analysis allowed us to compare the deliberate head motions carried out in our study participants with non-deliberate head motion that may be encountered in a typical clinical setting.

### 2.3 CNN-based head pose estimation

Video frames were first converted into x and y gradient images using a Sobel filter to reduce background noise. The input of the model at time *t_n_* was a concatenation of the gradient images obtained using: < *f_n_* − *f*_0_, *f*_*n*-1_ − *f*_0_, *f_n_* − *f*_*n*-1_ >, where *f_n_*, *f*_*n*-1_, *f*_0_ denote the x and y gradient images at time *t_n_*, *t*_*n*-1_, and *t*_0_ respectively. Our CNN model was based on the VGGNet network architecture [23]. We reduced the sizes of some convolutional and dense layers in the original VGG-16 model to protect against overfitting. We further added model regularization using a 0.5 dropout rate on the dense layers. We used our own initialization for the predictive model; no pre-training was used. Models were developed using the Keras API [24].

In our model architecture (Table 1), all filters were 3×3 and all convolutional layers had the same padding. The first two layers have 64 channels. Then after a max pooling layer of stride (2, 2), there are 2 additional convolutional layers with 128 filters. After another max pooling layer, there are 2 sets of 3 convolutional layers and a max pooling layer in between; each set has 256 and 512 filters respectively. Finally, there are four dense layers of size 512, 256, 128, 64 respectively.

Because most of the target rotation and translation values are close to 0, a standard loss function (e.g. mean squared error, MSE) would minimize the global loss, thereby training the model to underestimate large head motions. To address this issue, we adopted a cost sensitive learning approach [25]. We customized the model’s loss function to incorporate a weight proportional to the magnitude of the head move at each data point. Specifically, the training loss was defined as:

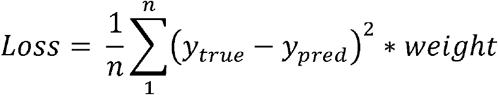

where *y_true_* is the ground-truth, *y_pred_* is the model’s prediction, and the *weight* is defined as:

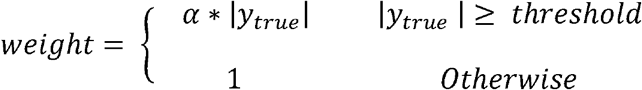

The *threshold* was set at 0.05 and 1.0 for the rotation and translation parameters respectively. The scaling constant α was set at 200 and 10 using the training data for the rotation and translation models respectively. Thus, the loss of data points whose head motions are above the *threshold* was weighted α ⁎ |*y_true_*| times over those whose head motions were below the *threshold*.

Two head pose prediction models were trained using the described approach. The first assessed the ability of our model to predict head pose in out-of-sample cases. For this model 14 subjects were used to train the model, and the model was used to estimate head pose changes from video obtained during EPI-BOLD acquisition in the remaining 7 hold-out subjects. Associated ground truth head pose estimates from the EPI-BOLD acquisition were used to assess model performance. Agreement between the video-based and EPI-BOLD head pose estimates over the duration of the EPI-BOLD acquisition was assessed by measuring the coefficient of determination (R^2^) and associated p-value for a linear model for each of the 6 head pose parameters comprising rotation around the x y and z planes and translation in each of the x y and z directions. These analyses consisted of 42 univariate analyses = 7 hold-out subjects × 6 head pose parameters; a Bonferroni corrected p-value = 0.05/42 = 1.2 × 10^−3^ was used.

The second head pose prediction model was trained on video obtained during EPI-BOLD acquisition in all 21 subjects. This model was used to directly estimate head pose changes using video obtained during structural scans in the study participants. Whole brain cortical thickness maps and volumes of subcortical neuroanatomical structures were estimated using Freesurfer v6.0 (https://surfer.nmr.mgh.harvard.edu/, [26, 27]). Because four structural MRI scans were available from each participant (two motion-affected and two motion-free scans), we utilized the longitudinal Freesurfer processing stream to improve within subject coregistration [28]. We compared video-based average head pose estimates obtained during motion-free structural scans with estimates obtained during motion-affected scans. We expected mean absolute z-axis rotation to be increased in the motion-affected scans, since the instructed head motion (“no”) during the scans is a rotation around this axis. The relationship between vertex-based cortical thickness estimates and video-based head pose estimates was investigated using a mixed linear model [29], with vertex-wise cortical thickness as the outcome variable, 7 fixed effects predictors consisting of the 6 average absolute head pose estimates: rotation and translation in the x,y and z planes (rot_x/y/z_ in radians and trans_x/y/z_ in mm), and age at time of scan. Each subject was modelled as a random effect. A mixed linear model allowed us to incorporate morphometric estimates derived from multiple scans of individual subjects with varying amounts of head motion. We derived vertex-wise maps of the estimated effect of z-axis rotation on cortical thickness in units of mm/radian across subjects using the standard Freesurfer *fsaverage* template. Similar analyses were undertaken with subcortical volume estimates obtained using Freesurfer. For these analyses, subcortical volume was the outcome variable with the six head pose parameters and age at time as scan modelled as fixed effects and subject modelled as a random effect.

## 3. RESULTS

Comparison of video-based head pose estimates with ground truth fMRI-based estimates in our hold-out dataset supports the utility of our proposed approach for estimating in-scanner head pose changes. An example comparison of video-based and fMRI-based head pose estimates in a participant from the hold-out dataset is provided in Figure 2. The video-based technique is primarily sensitive to rotational motion changes, which is to be expected given that the deliberate in-scanner head movements used to train the predictive model were rotational motions about the cardinal axes in subject space. Plots of the loss evolution as a function of the number of epochs are provided as supplementary material.

**Figure 2.**
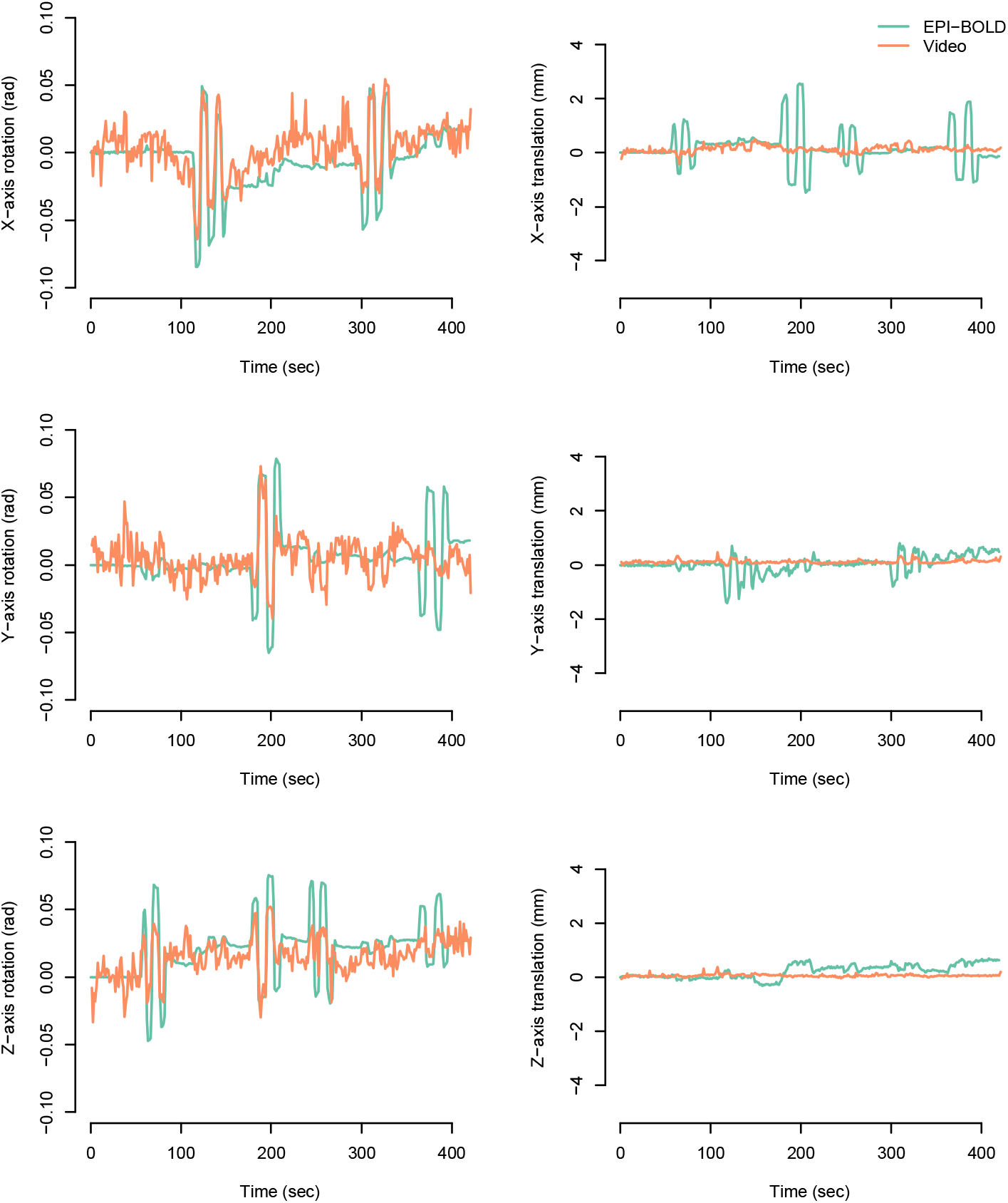
Head pose changes derived from in-scanner video (orange line) compared with head pose changes estimated using co-registration of sequential EPI volumes (green line) in a hold-out subject undertaking deliberate in-scanner head movements.

There was a statistically significant relationship between video-based head pose and fMRI-based head pose estimates in 5/7 subjects for rotation in the x-axis (median R^2^ = 0.34, p = 6.6 × 10^−31^), 6/7 subjects for rotation in the y axis (median R^2^ = 0.11, p = 6.2 × 10^−10^) and 6/7 subjects for rotation in the z-axis (median R^2^ = 0.21, p = 2.6 × 10^−18^). For video-based assessment of translational head pose changes, statistically significant correlations between video- and fMRI-based estimates were observed in 6/7 subjects for translation in the x-axis (median R^2^ = 0.29, p = 1.5 × 10^−25^), 4/7 cases for y-axis translation (median R^2^ = 0.06, p = 5.5 × 10^−6^) and 5/7 cases for z-axis translation (median R^2^ = 0.04, p = 1.5 × 10^−25^). The R^2^ distribution values are shown in Figure 3.

**Figure 3.**
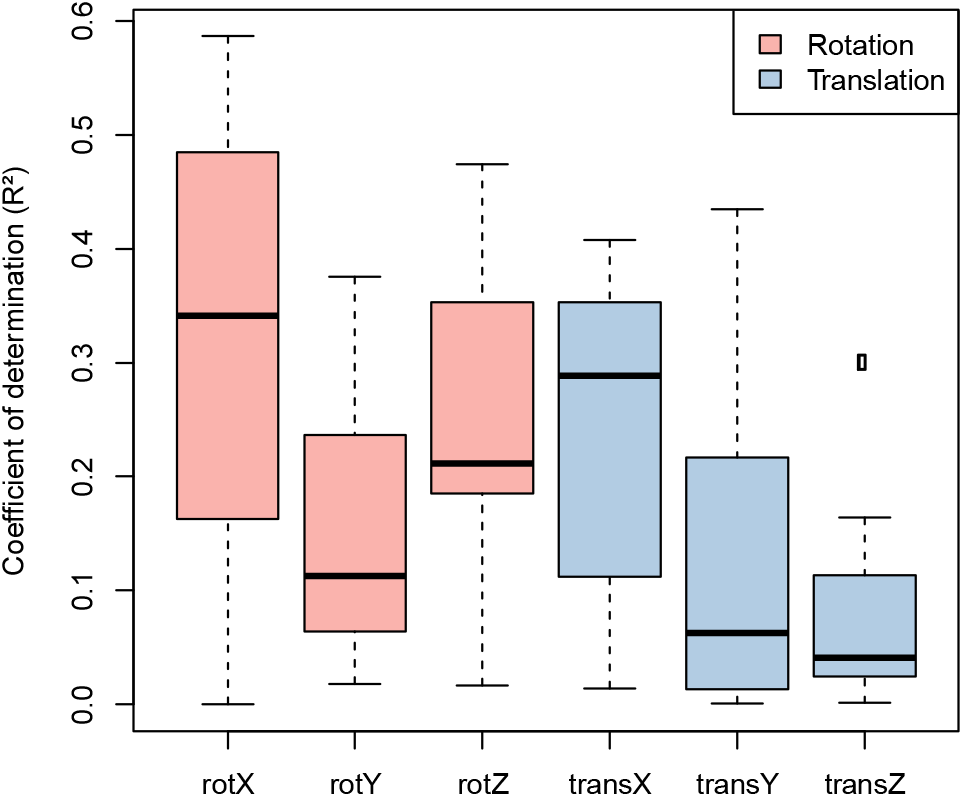
Distribution of R^2^ values indicating quality of fit between video-based head pose estimates and fMRI-based head pose estimates across all subjects in the hold-out dataset (N = 7). The figure shows that the ability of our technique to model head pose varies depending on the direction of head motion relative to the camera.

Video-based head pose changes obtained during motion-affected and motion-free structural MRI scans support the utility of our technique. For motion-affected structural MRI scans participants were instructed to only carry out “no” head motions, corresponding to rotation around the z-axis. Comparison of mean absolute values between motion-affected and motion-free scans for each of the six motion parameters identified significantly increased video-based estimates of z-axis rotation (7.6 × 10^−3^ radians, 99.2% confidence interval = [3.3 × 10^−3^, 1.2 × 10^−^ 2]) and x-axis translation (0.1 mm, 99.2% confidence interval = [0.04,0.17]) in motion affected scans relative to motion free scans (Figure 4). Note that a 99.2% confidence interval corresponding to a Bonferroni-corrected alpha = 0.05/6 was used. As expected, no statistically significant differences were identified for rotation around the x and y axes during the structural MRI scans.

**Figure 4.**
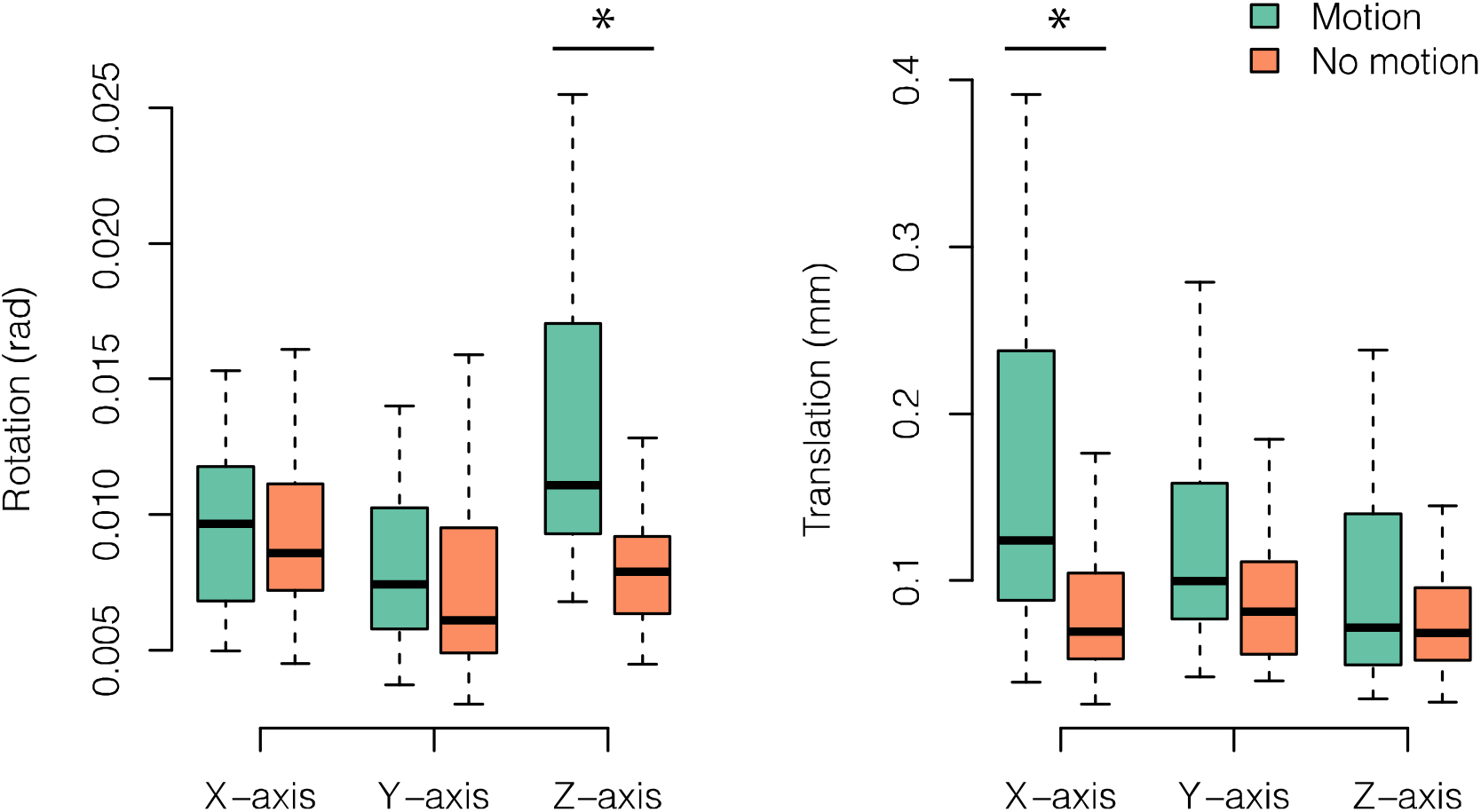
Video-based head pose estimates obtained during structural MRI acquisition in all study participants (N = 21). The boxplots show average values over the duration of the acquisition for motion-affected (green) and motion-free (orange) T1w acquisitions. The plots show significantly increased average head motion in z-axis rotation and x-axis translation (marked with asterisks), indicating that our video-based predictive model is sensitive to in-scanner head motion.

Estimation of the vertex-wise relationship between cortical thickness and average absolute rotational head motion around the z-axis indicates that increased head motion is associated with reduced cortical thickness over most of the cortex (Figure 5). For some regions, including bilateral anterior frontal lobes, central sulcus and occipital lobe, the opposite relationship was observed (i.e., increased head motion associated with increased cortical thickness). This spatially variable pattern is highly similar to a previous study by our group in an independent cohort [2]. For subcortical volume estimates a similar relationship was observed, with the predominant finding associated with increased head motion being reduced subcortical volume in 10/21 structures (Figure 6). For the remaining structures, no significant difference was observed in 7/21 structures, and increased volume with increased head motion was observed in four structures, left and right choroid plexus and vessel labels. Notably, neither the choroid plexus nor vessel labels are gray matter structures.

**Figure 5.**
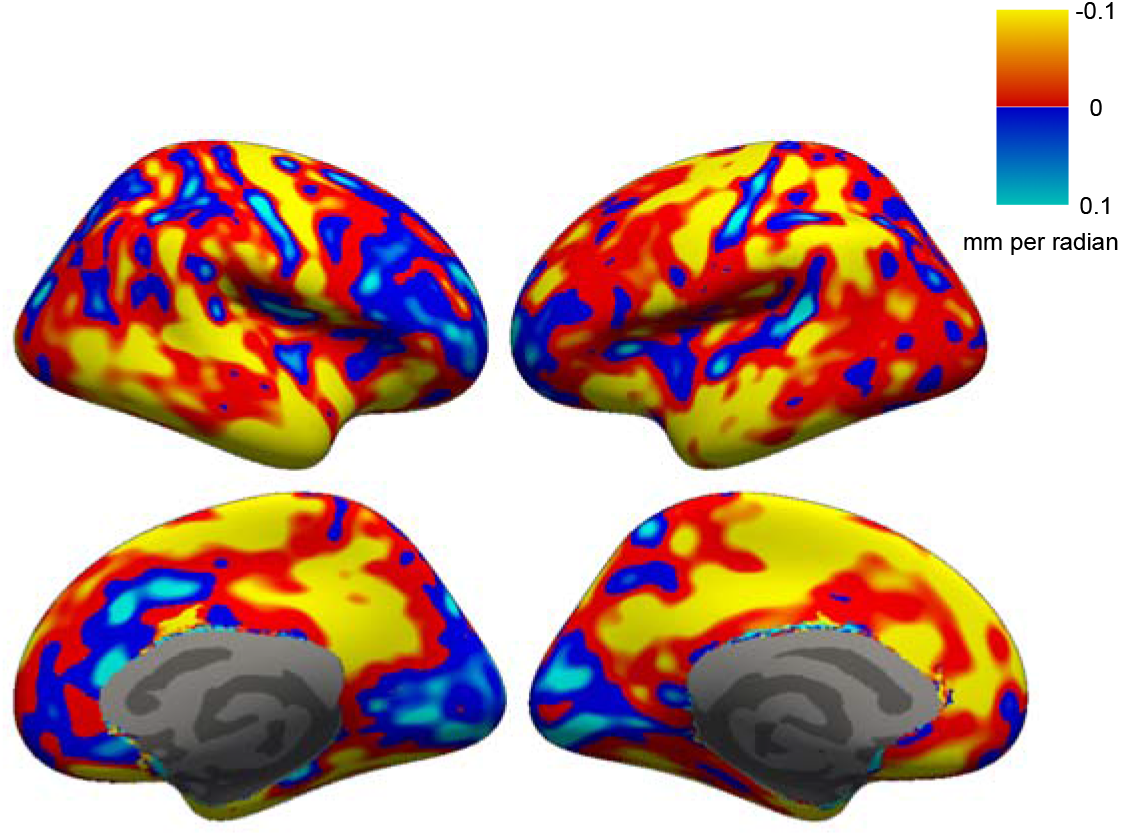
Relationship between video-based in-scanner head pose estimates and cortical thickness in study participants. The brain maps show the relationship between z-axis rotation and cortical thickness, with “hot” colors indicating regions where increased head motion is associated with decreased cortical thickness. The figure demonstrates that in-scanner head motion is associated with reduced cortical thickness changes over most of the cortex, with some regions (motor strip, occipital lobe and some frontal regions) showing the opposite relationship.

**Figure 6.**
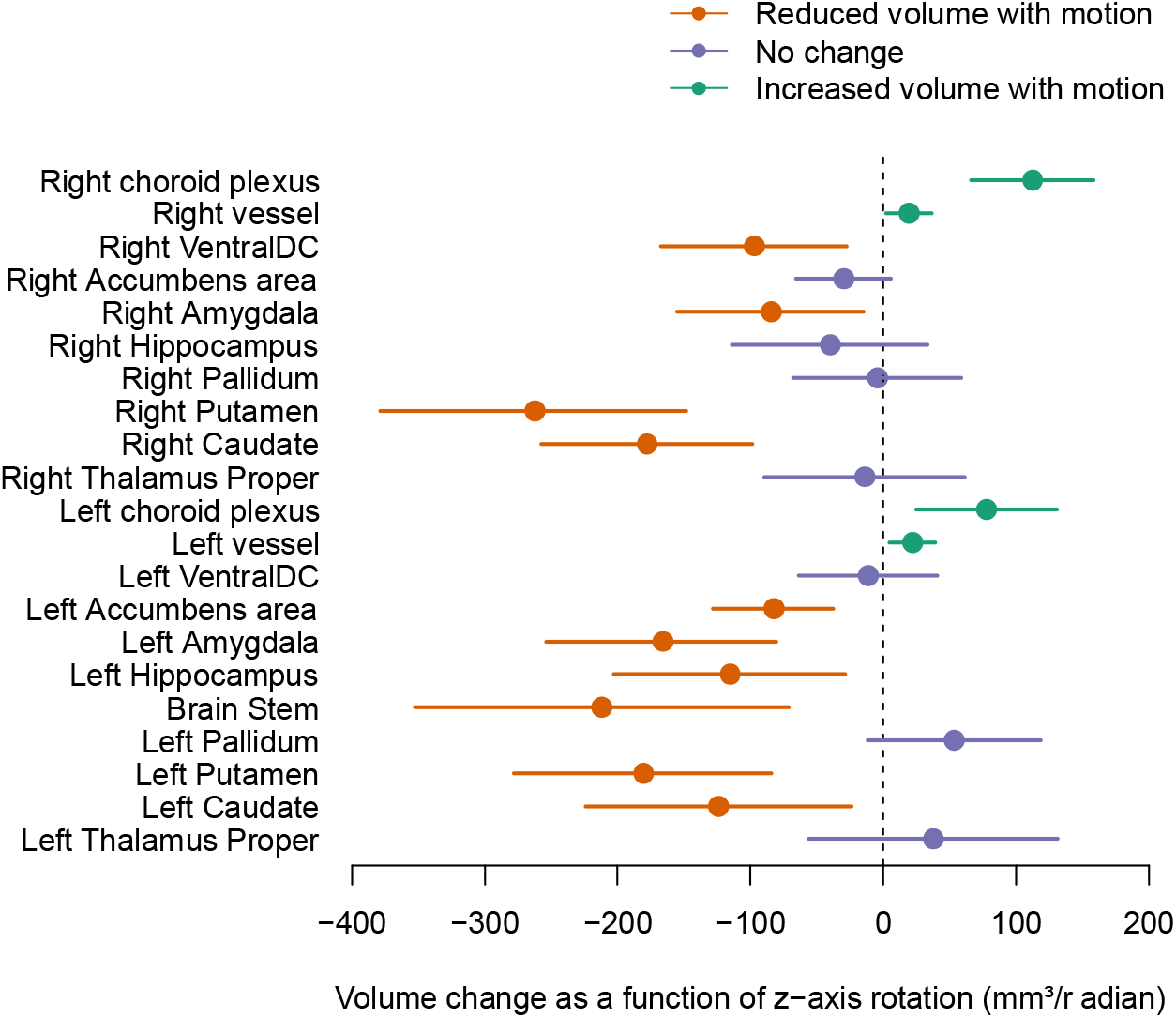
Increased in-scanner head motion is associated with changes in regional subcortical volume estimates. The extremes of the horizontal lines indicate the 95% confidence intervals. Increased head motion is associated with reduced volume in the hippocampus, amygdala and deep gray matter structures (orange lines). Purple lines indicate structures where no significant effects were observed, and increased volume with increased motion was observed in bilateral choroid plexus and vessel labels (purple lines).

## 4. DISCUSSION

We present a method to directly estimate changes in head pose during structural MRI scans. We trained a convolutional neural network model to estimate changes in head pose using video obtained from an in-scanner eye tracker. The CNN was trained using ground truth head pose estimates from an EPI-BOLD acquisition with participants carrying out deliberate in-scanner head motions. The predictive model was then used to estimate changes in head pose during acquisition of structural MRI scans. These video-based head pose estimates were used to statistically model the relationship between in-scanner head motion and quantitative estimates of cortical thickness and volume of subcortical brain structures. The cortical thickness and subcortical volume analyses demonstrated that there is a statistically significant relationship between in-scanner head motion and estimated morphometric properties, suggesting that our video-based technique will be useful to statistically control for the effects of head motion in clinical research studies that use structural MRI. One interesting observation is the remarkable similarity between cortical surface maps that show the vertex-wise distribution of the relationship between head motion and cortical thickness from this study (N = 21 subjects) and a larger study from our group that used proxy estimates of head motion derived from resting state fMRI acquisitions obtained in the same imaging session as the whole brain T1-weighted MRI scan (N = 2141 subjects, [2]). For both studies we observed a pattern of generally brain-wide reduced cortical thickness with increased head motion, with small regions of increased cortical thickness with increased head motion in the postcentral gyrus and occipital lobe. These latter regions correspond to brain regions with reduced cortical thickness and contrast between the cortical gray matter and underlying white matter. Although the precise mechanism underlying the variable motion-related changes across the cortex is unclear, it is likely that motion-related blurring interferes with delineation of the cortex/WM boundary, with blurring in low contrast regions skewing the cortical thickness estimates upwards, and the opposite effect occurring in high thickness/contrast regions.

In-scanner eye trackers are used in many clinical research imaging centers. Our technique has minimal technical requirements; for example, it does not require any markers to be attached to the participant, which means it may be readily implemented in both research and clinical imaging environments. Although we have demonstrated the utility of the technique for controlling for the effects of motion in cortical thickness and subcortical volume analyses, the method is independent of image acquisition protocol and could therefore also be useful for other MRI modalities in which head motion is an issue, such as diffusion imaging [30]. Our technique may also be useful for validating proposed proxy measures of in-scanner head motion that are derived from analysis of the structural MRI scans such as average edge strength [31, 32] or Euler number [33].

Although our evidence supports video-based predictive modeling approach to estimate and statistically control for head motion in MRI-based quantitative morphometry studies, several issues should be addressed before the technique is employed in clinical studies. Our participants deliberately moved their heads in the scanner. These head motions likely only model severe in-scanner head motions that may occur in a natural setting. Future work may be necessary to obtain training data to optimize performance in clinical imaging studies. Given the large number of studies that obtain both functional MRI data and structural MRI scans in a single imaging session, one potential avenue to explore would be to utilize a ‘big data’ approach to train a predictive model using video obtained during fMRI scanning sessions in clinical research participants. This would improve similarity between the type of head motions used to train the model and those encountered in natural settings, however it is likely that this approach would require a substantially larger number of subjects than those used in our study since large head motions are less common in a natural setting. In addition to increasing the number of subjects used to train the model, increasing the length of the EPI acquisition or shortening the repetition time to increase the sampling frequency of the training data is likely to yield improvements in model performance. The 7 minute acquisition time in our study was chosen to allow enough time to carry out the three types of head motion in our study, along with 2 minutes of motion-free acquisition time. The relationship between the EPI acquisition time and model performance is currently unclear but would likely improve with longer acquisitions.

Another limitation of our work is the relatively modest size of our dataset. Thus, generalizability of our model is unclear and may be limited. For example, the scanner operator often modifies the positioning of the eye tracker when preparing a participant for scanning, which may lead to variable positioning of the face relative to the eye tracker camera. Our study did not investigate how this source of variability affects model performance. Alternative approaches such as landmark-based methods may prove to be useful for addressing this issue [34]. Also, we used the average head motion for each cardinal axis over the duration of the scan as predictor variables for our morphometric analyses. There may be improved metrics that could be extracted from the time series head motion data to use as explanatory variables; for example, Afakan et al. found that motion free time as a proportion of the total acquisition time was highly correlated with image quality in a prior study investigating head motion during MRI in pediatric subjects [15]. Finally, given that deep learning is a rapidly evolving field, future methodological developments in deep learning techniques may improve model performance. For this reason, we have made imaging and video data from this study available to the neuroscientific community to facilitate development of alternative approaches.

In summary, we have presented a video-based deep learning technique for directly estimating changes in head pose during structural MRI acquisitions. We have demonstrated that our approach can be used to estimate and statistically control for the effects of head motion on morphometric estimates derived from structural MRI data.

## Acknowledgements

This work was supported by FACES foundation, NY, USA. The authors thank S. Robert Frost for their helpful comments on an earlier version of the manuscript.

## REFERENCES

1. Reuter, M., et al., Head motion during MRI acquisition reduces gray matter volume and thickness estimates. Neuroimage, 2015. 107: p. 107–115.

2. Pardoe, H.R., R. Kucharsky Hiess, and R. Kuzniecky, Motion and morphometry in clinical and nonclinical populations. Neuroimage, 2016. 135: p. 177–85.

3. Alexander-Bloch, A., et al., Subtle in-scanner motion biases automated measurement of brain anatomy from in vivo MRI. Hum Brain Mapp, 2016. 37(7): p. 2385–97.

4. Savalia, N.K., et al., Motion-related artifacts in structural brain images revealed with independent estimates of in-scanner head motion. Hum Brain Mapp, 2017. 38(1): p. 472–492.

5. Tisdall, M.D., et al., Prospective motion correction with volumetric navigators (vNavs) reduces the bias and variance in brain morphometry induced by subject motion. Neuroimage, 2016. 127: p. 11–22.

6. Murphy-Chutorian, E. and M.M. Trivedi, Head pose estimation in computer vision: a survey. IEEE Trans Pattern Anal Mach Intell, 2009. 31(4): p. 607–26.

7. Patacchiola, M. and A. Cangelosi, Head pose estimation in the wild using Convolutional Neural Networks and adaptive gradient methods. Pattern Recognition, 2017. 71: p. 132–143.

8. Maclaren, J., et al., Prospective motion correction in brain imaging: a review. Magn Reson Med, 2013. 69(3): p. 621–36.

9. Godenschweger, F., et al., Motion correction in MRI of the brain. Phys Med Biol, 2016. 61(5): p. R32–56.

10. White, N., et al., PROMO: Real-time prospective motion correction in MRI using image-based tracking. Magn Reson Med, 2010. 63(1): p. 91–105.

11. Kyme, A.Z., et al., Marker-free optical stereo motion tracking for in-bore MRI and PET-MRI application. Med Phys, 2020. 47(8): p. 3321–3331.

12. Slipsager, J.M., et al., Markerless motion tracking and correction for PET, MRI, and simultaneous PET/MRI. PLoS One, 2019. 14(4): p. e0215524.

13. Gholipour, A., et al., Motion-Robust MRI through Real-Time Motion Tracking and Retrospective Super-Resolution Volume Reconstruction. 2011 Annual International Conference of the Ieee Engineering in Medicine and Biology Society (Embc), 2011: p. 5722–5725.

14. Kober, T., et al., Head Motion Detection Using FID Navigators. Magnetic Resonance in Medicine, 2011. 66(1): p. 135–143.

15. Afacan, O., et al., Evaluation of motion and its effect on brain magnetic resonance image quality in children. Pediatr Radiol, 2016. 46(12): p. 1728–1735.

16. Maclaren, J., et al., Measurement and correction of microscopic head motion during magnetic resonance imaging of the brain. PLoS One, 2012. 7(11): p. e48088.

17. Tisdall, M.D., et al., Volumetric navigators for prospective motion correction and selective reacquisition in neuroanatomical MRI. Magn Reson Med, 2012. 68(2): p. 389–99.

18. Frost, R., et al., Markerless high-frequency prospective motion correction for neuroanatomical MRI. Magn Reson Med, 2019. 82(1): p. 126–144.

19. Harms, M.P., et al., Extending the Human Connectome Project across ages: Imaging protocols for the Lifespan Development and Aging projects. Neuroimage, 2018. 183: p. 972–984.

20. Casey, B.J., et al., The Adolescent Brain Cognitive Development (ABCD) study: Imaging acquisition across 21 sites. Dev Cogn Neurosci, 2018. 32: p. 43–54.

21. Jenkinson, M., et al., Improved Optimization for the Robust and Accurate Linear Registration and Motion Correction of Brain Images. NeuroImage, 2002. 17(2): p. 825–841.

22. Di Martino, A., et al., The autism brain imaging data exchange: towards a large-scale evaluation of the intrinsic brain architecture in autism. Mol Psychiatry, 2014. 19(6): p. 659–67.

23. Simonyan, K. and A. Zisserman Very Deep Convolutional Neural Networks for Large-Scale Image Recognition. arXiv, 2015.

24. Chollet, F., Keras. 2015.

25. Fernández, A., et al., Cost-sensitive learning, in Learning from Imbalanced Data Sets. 2018, Springer. p. 63–78.

26. Fischl, B. and A.M. Dale, Measuring the thickness of the human cerebral cortex from magnetic resonance images. Proc Natl Acad Sci U S A, 2000. 97(20): p. 11050–5.

27. Fischl, B., et al., Whole brain segmentation: automated labeling of neuroanatomical structures in the human brain. Neuron, 2002. 33(3): p. 341–55.

28. Reuter, M., et al., Within-subject template estimation for unbiased longitudinal image analysis. Neuroimage, 2012. 61(4): p. 1402–18.

29. Bates, D., et al., Fitting Linear Mixed-Effects Models Using lme4. Journal of Statistical Software; Vol 1, Issue 1 (2015), 2015.

30. Baum, G.L., et al., The impact of in-scanner head motion on structural connectivity derived from diffusion MRI. Neuroimage, 2018. 173: p. 275–286.

31. Aksoy, M., et al., Hybrid prospective and retrospective head motion correction to mitigate cross-calibration errors. Magn Reson Med, 2012. 67(5): p. 1237–51.

32. Zaca, D., et al., Method for retrospective estimation of natural head movement during structural MRI. J Magn Reson Imaging, 2018. 48(4): p. 927–937.

33. Rosen, A.F.G., et al., Quantitative assessment of structural image quality. Neuroimage, 2018. 169: p. 407–418.

34. Smith, B.M., et al., Nonparametric Context Modeling of Local Appearance for Pose- and Expression-Robust Facial Landmark Localization. 2014 Ieee Conference on Computer Vision and Pattern Recognition (Cvpr), 2014: p. 1741–1748.

