## Supplemental Data for "Estimation of in-scanner head pose changes during structural MRI using a convolutional neural network trained on eye tracker video"

The following figures show the evolution of model loss for training and testing datasets as a function of the number of epochs. The figures show results for the model trained on N=14 subjects.

| 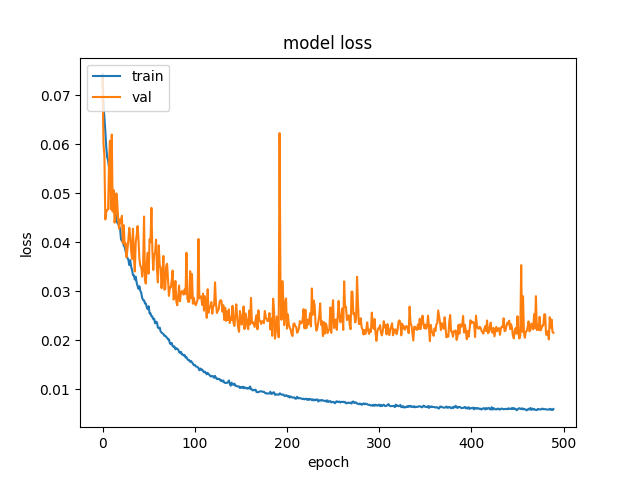 | 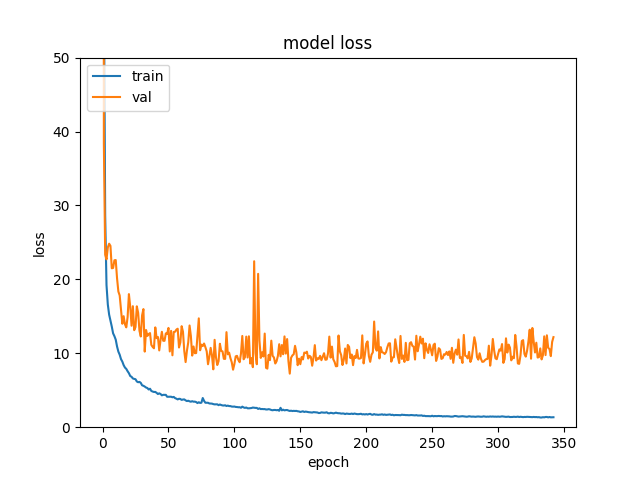 |
| --- | --- |
| Figure 1. The evolution of model loss for training and testing datasets as a function of the number of epochs. The figures show results for the model trained on N=14 subjects. The left figure shows data for rotation predictions, and the right shows data for translation predictions. | |
